# Structome-AlignViewer: On Confidence Assessment in Structure-Aware Alignments

**DOI:** 10.1101/2025.05.31.657027

**Authors:** Ashar J. Malik, Siying Mao, Philip Hugenholtz, David B. Ascher

## Abstract

Protein structure-based comparison provides a framework for uncovering deep evolutionary relationships that can escape conventional sequence-based approaches. Encoding three-dimensional protein structures using a simplified structure-aware alphabet, can lead to compact, comparable strings that retain key spatial relationships. Although this enables comparison, structure-aware alignments can experience misaligned regions, particularly when comparing proteins with substantial divergence in fold architecture. To address this, a web-based resource, Structome-AlignViewer, is introduced in this work for evaluating the quality of structure-aware alignments through both spatial mapping of alignment columns to protein structures and quantitative confidence scoring. Confidence is computed from pairwise structural substitutions between adjacent inputs and normalized within each alignment to highlight relatively well-supported columns. To provide broader context, thousands of alignments from established structural classification systems were analysed, allowing for an empirical comparative statistic to be derived to assess alignment quality. Option to exclude gap-rich regions enable users to refine alignments and focus on conserved structural cores. This approach provides an interpretable method for assessing structural alignment quality and supports more robust comparative and evolutionary analyses. Structome-AlignViewer is freely available at https://biosig.lab.uq.edu.au/structome_alignviewer/.

## Introduction

Determining homology among biological macromolecules has long been central to unravelling the history of life on Earth. By identifying relationships among genes and proteins, homology inference provides the scaffolding upon which evolutionary biology, comparative genomics, and structural bioinformatics are built.

Traditionally, molecular sequence data have served as the primary source for detecting evolutionary relationships [1]. While comparative sequence-based analysis has driven decades of discovery, it faces a fundamental limitation: as sequences diverge over deep evolutionary time, the homology signal progressively erodes, making it increasingly difficult to discern signal from noise. This loss of detectable similarity defines the so-called “twilight zone” [2] of sequence comparison, beyond which many evolutionary relationships are missed.

Protein structure, in contrast, tends to be more robust to changes in the underlying amino acid sequence [3, 4, 5]. It preserves key aspects of molecular function and folding architecture even as mutations accumulate and sequence identity diminishes. This robustness has made structural data an appealing alternative for recovering deeper evolutionary signals [5, 6]. A recent advance in defining a structural alphabet—the introduction of 3Di states used by Foldseek [7]—has opened a new frontier. By converting protein structures into 3Di sequences, classical alignment and phylogenetic tools can be applied to this structure-aware alphabet, effectively extending the temporal reach of evolutionary inference [8, 9].

This structure-informed evolutionary analysis generally follows a three-step process [8]: (1) compile a dataset of proteins [10, 11], (2) align their structure-aware sequences, and (3) build a phylogenetic tree from the resulting alignment [8]. As promising as this approach is, the novelty of the method raises important open questions about how best to apply and interpret the inference.

One key aspect, of the many, that warrants a deeper exploration is the quality of the alignment itself where even slight column misplacements can distort phylogenetic inference by skewing comparative estimates and resulting tree topologies [12, 13, 14]. While the structure-aware alignment strategy successfully incorporates three-dimensional context, at present users lack a straightforward way to (i) visually map regions back to the actual structures for manual review and (ii) assess per-column reliability. Consequently, it can be difficult to refine this new class of alignments and ensure that downstream maximum likelihood or other statistical phylogenetic methods receive correct inputs.

In this work, Structome-AlignViewer, an interactive, web-based platform is introduced that addresses this challenge by providing a fast and user-friendly environment to compute, inspect, and assess the quality of structure-aware alignments. The platform not only links alignment columns directly to 3D structures through an interactive and user-friendly interface, but also incorporates a statistical framework to quantify both per-column and overall alignment confidence. To contextualise these confidence values, an empirical alignment analysis across structurally homologous groups defined in SCOP [15, 16] and CATH [17] was carried out, yielding a background distribution of confidence scores. This distribution serves as a reference for assessing the significance of average confidence values in user-generated alignments.

Overall, this resource is intended to provide more transparency for users carrying out structure-based phylogenetic analysis, and hence is intended to be an important advance in this area.

## Methods

### Structure-aware Sequence Generation

This work used Foldseek to generate structure-aware sequences of proteins using their respective 3D structures. Foldseek assigns each amino acid in the protein structure a structure-aware character based on the tertiary interactions made by the respective residue. This results in a sequence of structure-aware characters, which collectively are referred to as a structure-aware sequence in this work.

### Aligning Structure-aware Sequences

The alignment of structure-aware sequences was carried out using ClustalW2 [18], applying a conventional multiple sequence alignment algorithm to the structure-aware representation. A custom substitution matrix [7], included with Foldseek, was provided as input to ClustalW2, which is necessary since a distinction must be made between amino acids and structure-aware characters. Furthermore, alignments were performed in the “slow” mode to maximize alignment quality.

### Per-Column Confidence Metric

To assess alignment quality, a per-column confidence score *C*_*j*_ is computed for each alignment column *j* based on the sum of substitution scores between adjacent sequences in the alignment (see supplementary data for examples). For each column, all adjacent sequence pairs are considered:

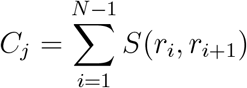

where:

- *N* is the number of sequences in the alignment,
- *r*_*i*_ and *r*_*i*+1_ are the structure-aware characters from adjacent sequences at column *j*,
- S(*r*_*i*_,*r*_*i*+1_) is the substitution score for that pair, obtained from the Foldseek substitution matrix.

Adjacent pairs involving gaps are excluded from analysis. The resulting raw column scores capture local structural coherence among neighbouring sequences.

To normalize these values within each alignment, min-max scaling is applied across all columns:

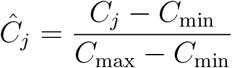

where *C*_min_ and *C*_max_ are the minimum and maximum raw confidence scores across all columns in the alignment. The resulting normalized score 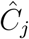 lies in the range [0, 1] and reflects the strength of structural agreement for each column relative to the weakest and strongest-aligned positions in the same alignment.

An average of all normalized column scores is also computed per alignment:

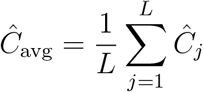

where *L* is the total number of columns. This serves as a representative measure of overall alignment confidence.

### Interpretation of the Confidence Metric

To interpret confidence values statistically, thousands of alignments of SCOP and CATH groupings were used. These alignments have previously been used in other work [9]. Briefly, at the family level in SCOP and the homology level in CATH, protein structures were retrieved, and domain regions were extracted based on annotations from the respective classification databases. As described above, these structures were converted into structure-aware sequences and alignments were generated using the ClustalW2 method.

Two thresholds were applied to filter alignments for analysis:

- alignments comprising *≥*10 structures, and
- alignments of length *≥*100 residues.

This resulted in 2,089 SCOP alignments and 1,931 CATH alignments. Confidence scores were calculated for each alignment, and the corresponding means and standard deviations were determined for both classification systems.

Z-scores reported on the results page on Structome-AlignViewer are derived from these distributions and computed separately for SCOP and CATH. The average of the two Z-scores 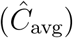 is used to assign a qualitative confidence category for each alignment (see Table 1). This approach offers users a way to interpret alignment quality in a statistically meaningful way.

**Table 1.** Interpretation of Z-score-based alignment confidence levels, based on statistical rarity.

| Z-score Range | Interpretation |
| --- | --- |
| $\hat{C}_{\text{avg}} \geq 2$ | Statistically Rare Alignment |
| $1 \leq \hat{C}_{\text{avg}} < 2$ | Above Average, Less Common |
| $-1 < \hat{C}_{\text{avg}} < 1$ | Common, Expected Range |
| $-2 < \hat{C}_{\text{avg}} \leq -1$ | Below Average, Less Common |
| $\hat{C}_{\text{avg}} \leq -2$ | Statistically Rare Alignment |

### Implementation of the Web Interface

The Structome-AlignViewer resource is implemented using Flask, Docker and Nginx. On the landing page of this resource, users can provide a delimited list of RCSB PDB accessions, including chain identifiers, which if valid are automatically converted into structure-aware sequences and subsequently aligned, and analysed. All structures are downloaded from RCSB PDB at runtime. Given compute limitations, a user is limited to 50 input structures per analysis.

The result page uses the Mol^*∗*^ [19] viewer to display protein structures. The feature viewer shows the alignment, in which columns are clickable, allowing users to instantly view the corresponding column in the structure selected. A drop-down menu allows users to toggle between different structures in the alignment. Furthermore, confidence scores are also encoded as B-factors in the rendered structures, enabling fast, intuitive colour-based visualization. Low confidence values approaching “0” appear towards the blue end of the spectrum and high confidence values approaching “1” appear towards the red end of the colour scale. Additionally users can obtain publication quality images from the native image export utility in Mol^*∗*^.

Structome-AlignViewer also implements a Gblocks-style trimming option [20], which removes alignment columns with *>*50% gap content. Raw structure-aware sequences for all structures along with both full and trimmed alignments and the dendrogram, which is also displayed on the results page, are available for download.

Biopython is used both on Structome-AlignViewer and for other analysis included in this work to parse and analyse alignments and load confidence values as B-factors in the respective protein structures for subsequent visualization in the Mol^*∗*^ viewer.

## Results

### Background Confidence Distributions from SCOP and CATH

To contextualize the confidence scores computed for user-submitted alignments, background confidence scores distributions from curated SCOP and CATH alignments were determined. These distributions serve as the statistical reference against which Z-scores are calculated. The violin plots shown in Figure 1 show these distributions for both SCOP and CATH datasets, comparing the full alignments (blue) to their trimmed counterparts (orange).

**Figure 1.**
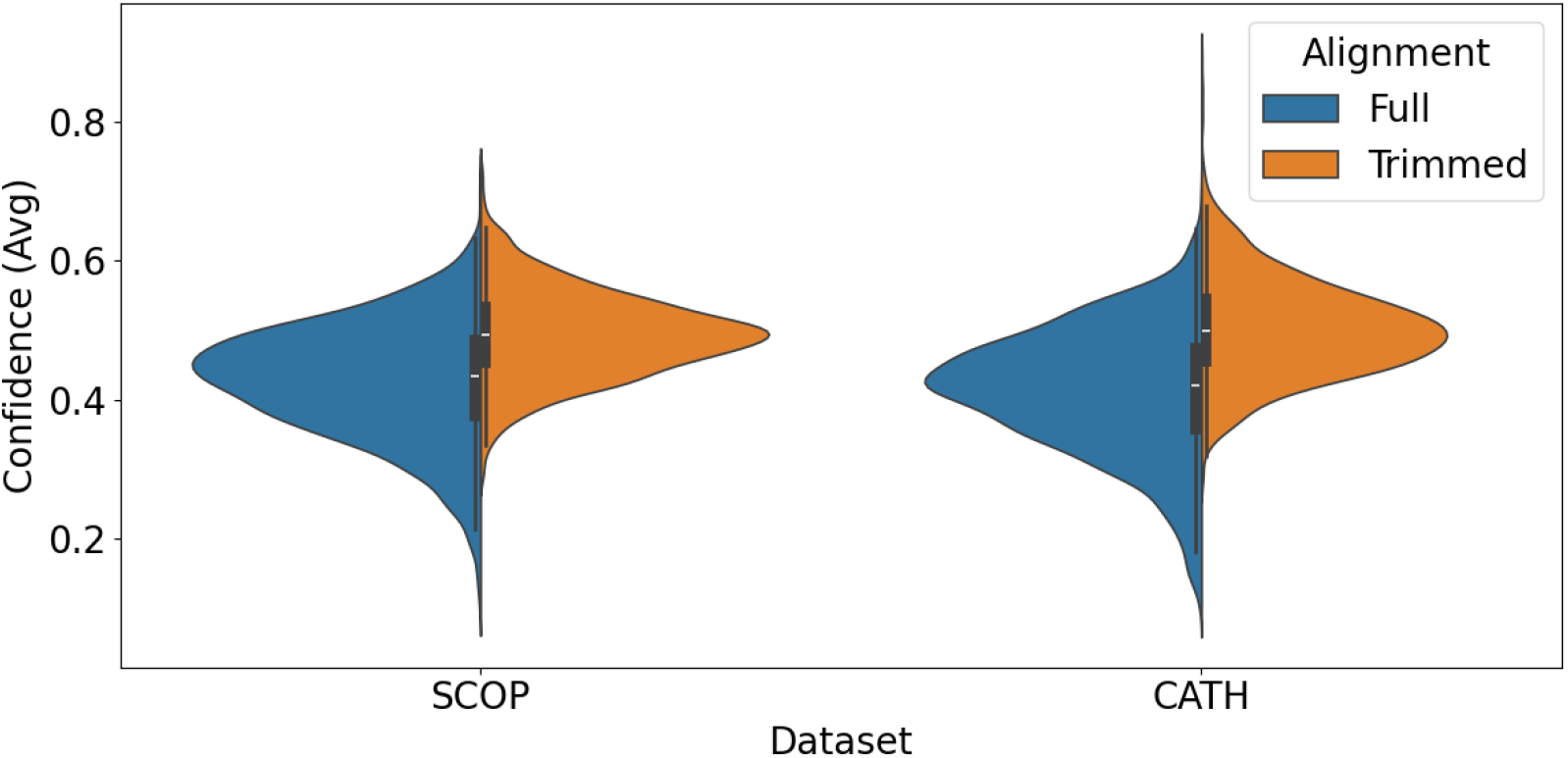
Distribution of average confidence scores for SCOP and CATH alignments. Each dataset is shown for both full alignments and their trimmed versions (Gblocks-style trimming with *>*50% gap removal).

As shown, trimming columns with high gap content results in a modest increase in average confidence since the gapped columns do not contribute to the score and the trimming results in reduced lengths, thus increasing the score. The trimmed alignments can therefore be interpreted as a refined subset, potentially more informative and suitable for downstream applications.

### Relationship Between Confidence and Gap Fraction

To further understand the drivers of alignment confidence, the relationship between gap fraction (i.e., proportion of gaps in the alignment) and the average confidence score was examined. As expected, a strong negative correlation was observed: alignments with higher gap content tend to exhibit lower average confidence. Figures 2 and 3 show this inverse relationship for SCOP and CATH alignments, respectively. These results reinforce the assumption that gaps often reflect regions of uncertainty or structural divergence. A high gap fraction indicates that many sites lack consistently aligned counterparts—whether due to flexible/disordered segments or failure to identify a common structural core. Such columns reduce alignment reliability and introduce noise in downstream analyses. Thus, removing gap-rich columns can reduce poorly supported positions and enrich the alignment.

**Figure 2.**
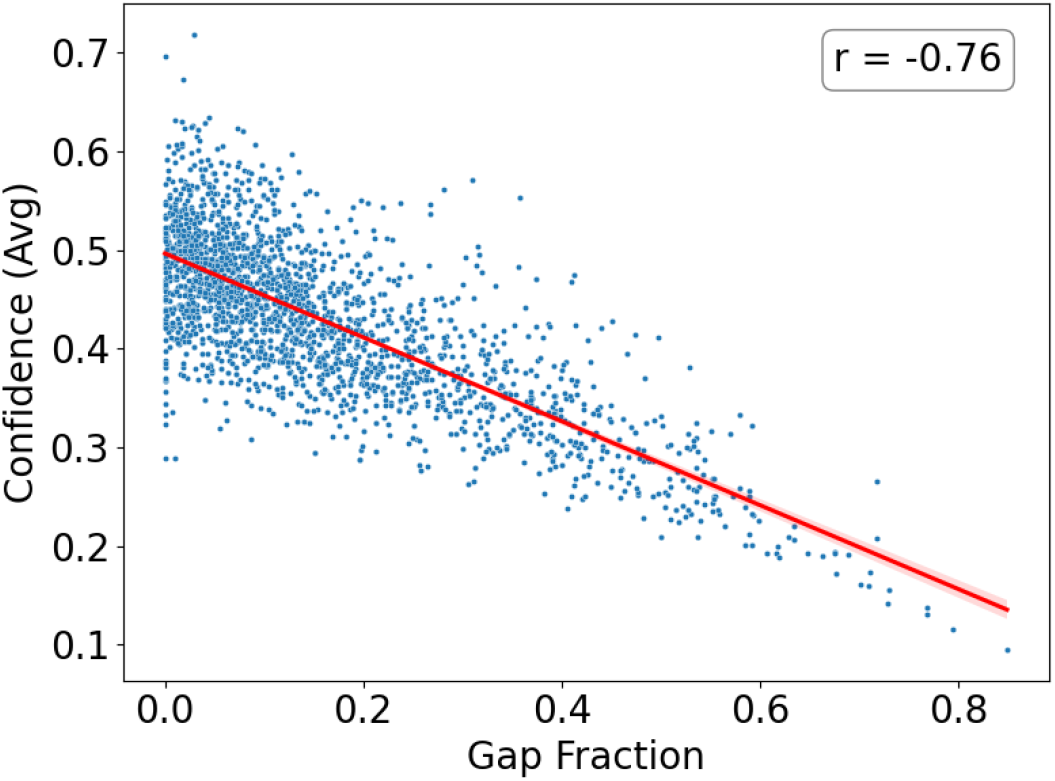
Correlation between gap fraction and average confidence score for SCOP alignments. Each point represents a single alignment, and a linear regression line (with 95% confidence interval) is overlaid in red. A strong negative correlation is observed (r=*−*0.76).

**Figure 3.**
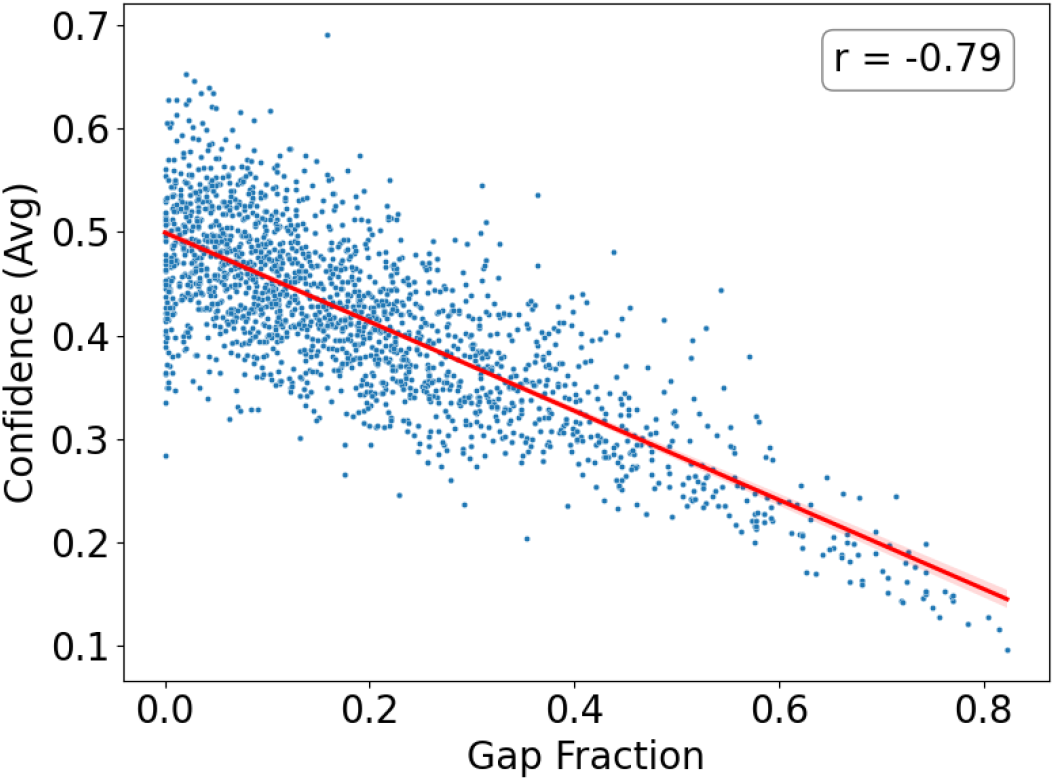
Correlation between gap fraction and average confidence score for CATH alignments. Each point represents a single alignment, and a linear regression line (with 95% confidence interval) is overlaid in red. Similar to SCOP, a strong negative correlation is observed (r=*−*0.79).

Building on this, the gap–confidence relationship also sets the stage for confidence-guided pruning strategies, in which columns with poor structural support or excessive gaps are systematically omitted. Refined alignments generated by this approach are well suited for non-parametric bootstrapping or any phylogenetic pipeline where reducing alignment noise is crucial. Thus, gap fraction emerges not merely as a technical artifact but as a biologically meaningful indicator of alignment quality—and, by extension, a practical filter for enhancing it.

## Discussion

### Assessing alignments using the confidence metric

Any structural alignment carried out for the purpose of determining homology, aims to identify equivalent positions across proteins compared. These equivalent positions then form the backbone of evolutionary comparison, delineating how folds are conserved, diverge, or adapt across the course of their respective evolutionary trajectories. Simply encoding 3D structures into structure-aware sequences before comparison does not necessarily guarantee a better alignment. As proteins diverge, segments may be introduced, which although part of a common fold may lack clear structural correspondence across the proteins, becoming especially pronounced when aligning distantly related proteins.

To determine if the alignment is reliable, the resource presented in this work, Structome-AlignViewer, offers two complementary layers of inspection. First, it enables users to directly map alignment columns back onto 3D structures which are rendered using the Mol^*∗*^ protein structure viewer. This spatial mapping helps assess whether incompatible structure-aware characters have been aligned. Secondly, the confidence score brings statistical context and provides a way to evaluate alignment quality against the background alignments generated from curated SCOP and CATH datasets.

The confidence metric derived per alignment originates from the same substitution matrix used during the alignment process. While the alignment aims to optimally determine overlap of structure-aware sequences, the confidence score adds an additional dimension, on a per-column basis, to the resulting alignment. While the scoring and alignment frameworks differ mathematically, they are unified in vocabulary and logic, making the confidence score a natural extension of the alignment itself.

Additionally, confidence values are scaled to highlight the best and worst aligned columns and thus reflect relative structural consistency rather than an absolute quality metric. Within each column, both identical and mismatched structure-aware characters are scored according to the structural alphabet’s substitution matrix. The normalization of all raw scores to a [0,1] range based on the alignment’s highest and lowest performing columns, not only captures whether residues agree but also how strongly they agree. As a result, columns that outperform the alignment’s average become visibly more trustworthy, highlighting where the alignment is particularly well supported.

Furthermore, to add context to user-generated structure-aware sequence alignments, the average normalized confidence score is computed and compared to the distributions of the same metric derived from thousands of SCOP and CATH alignments. Although each alignment is normalized independently, these averages form meaningful empirical distributions. The resulting Z-scores allow users to contextualise their alignment within the known homologous families. While this score does not predict phylogenetic correctness, it offers a principled and statistically grounded way to assess alignment quality, and can help guide decisions about trimming, weighting, or downstream analysis.

### Limitations of the confidence metric

It must be emphasized that alignment confidence is not equivalent to phylogenetic confidence. While a reliable alignment is essential for downstream tree construction, high alignment confidence does not imply correctness of the inferred topology. Many other factors—substitution model choice, rate heterogeneity, long-branch attraction, and sampling—can influence phylogenetic inference independently of alignment quality.

The purpose of Structome-AlignViewer is to provide users with a clearer view of how well their alignment represents conserved structural signals. Properly interpreted, confidence scores can guide alignment refinement without overextending their meaning whereas over-interpreting the score beyond its intended scope risks conflating alignment quality with evolutionary truth—a distinction that remains critical.

## Conclusion

Structome-AlignViewer, the resource introduced in this work, provides a robust framework for users to confidently assess structure-aware sequence alignments. It achieves this using two complementary methods. It allows visual inspection of alignment columns and additionally associates a confidence score with these columns. This combination allows users to assess the overall reliability of an alignment and identify regions that are well aligned across the protein structures compared. Additionally, to ensure consistency, the confidence score is calculated using the same substitution matrix employed for generating the alignment and is further normalised to facilitate comparisons both within a single alignment, and for computing averages which are then scored against extensive SCOP and CATH-derived distributions to provide valuable context.

While the confidence score, developed in this work, is not intended to be used towards assessing phylogenetic accuracy, it does open valuable avenues for further refinement. One immediate application is alignment pruning—removing poorly supported columns prior to tree inference. For example, in the case of non-parametric bootstrapping eliminating noisy positions could enhance signal-to-noise ratio and lead to reduced uncertainty in inferred trees.

As structure-based phylogenetics matures, Structome-AlignViewer will be instrumental in enhancing the reliability of structural comparisons carried out using structure-aware sequences and the evolutionary conclusions drawn from them.

## Availability

Structome-AlignViewer is freely available as a web application at: https://biosig.lab.uq.edu.au/structome_alignviewer/

