## Supplementary file for "Structome-AlignViewer: On Confidence Assessment in Structure-Aware Alignments"

### Supplementary example: Understanding confidence score using a toy example

To illustrate how per-column confidence scores are normalized within an alignment, a toy example involving 10 columns from a hypothetical structure-aware sequence alignment is used. Each column is scored based on the **sum of substitution scores between adjacent sequence pairs**, using the substitution matrix.

#### Raw Confidence Scores

The raw confidence score $C_j$ for each column is computed by summing the substitution scores between adjacent sequence pairs in that column. Let us assume the following raw scores:

| **Column** | **Raw Score** ($C_j$) |
|---|---|
| 1 | $-5.0$ |
| 2 | $-1.5$ |
| 3 | $-1.0$ |
| 4 | $-2.0$ |
| 5 | $-1.3$ |
| 6 | $-0.8$ |
| 7 | $-1.2$ |
| 8 | $-1.1$ |
| 9 | $-0.5$ |
| 10 | 0.0 |

Table 1: Raw confidence scores per column based on adjacent sequence pair substitution sums.

#### Min-Max Normalization

Min-max normalization is applied to rescale all scores between 0 and 1:

$$\hat{C}_j = \frac{C_j - C_{\min}}{C_{\max} - C_{\min}}$$

with:

$$C_{\min} = -5.0, \quad C_{\max} = 0.0$$

| Column | Normalized Score ($\hat{C}_j$) |
|---|---|
| 1 | 0.00 |
| 2 | 0.70 |
| 3 | 0.80 |
| 4 | 0.60 |
| 5 | 0.74 |
| 6 | 0.84 |
| 7 | 0.76 |
| 8 | 0.78 |
| 9 | 0.90 |
| 10 | 1.00 |

Table 2: Normalized confidence scores per column.

### Interpretation

This example highlights how min-max normalization allows confidence values to be interpreted in relative terms within the alignment. Although columns 2–10 vary in their raw scores, they all score considerably higher than column 1, which contains structurally incoherent residue pairs. Consequently, even moderately supported columns (e.g., column 4) receive high normalized scores (0.6), because they are much more coherent than the worst column.

- The lowest scoring column ($C_1$) is mapped to 0.
- The highest scoring column ($C_{10}$) is mapped to 1.
- All other columns are scaled between these two extremes.

This approach ensures that confidence scores are contextualized within the alignment, and motivates the use of empirical benchmarking (e.g., via SCOP and CATH) to interpret how strong a given alignment is compared to structurally curated families.

### Supplementary Example: Understanding Confidence Scores from a Real Alignment

To illustrate how the confidence scoring system works in practice, a real example alignment of five structures is used from which five aligned columns are drawn. Confidence scores are computed using pairwise substitution values drawn from the Foldseek substitution matrix. This matrix defines substitution preferences between 20 structural states and is used both during alignment and confidence evaluation.

The substitution matrix is shown below:

| | A | C | D | E | F | G | H | I | K | L | M | N | P | Q | R | S | T | V | W | Y | X |
|---|---|---|---|---|---|---|---|---|---|---|---|---|---|---|---|---|---|---|---|---|---|
| A | 6 | -3 | 1 | 2 | 3 | -2 | -2 | -7 | -3 | -3 | -10 | -5 | -1 | 1 | -4 | -7 | -5 | -6 | 0 | -2 | 0 |
| C | -3 | 6 | -2 | -8 | -5 | -4 | -4 | -12 | -13 | 1 | -14 | 0 | 0 | 1 | -1 | 0 | -8 | 1 | -7 | -9 | 0 |
| D | 1 | -2 | 4 | -3 | 0 | 1 | 1 | -3 | -5 | -4 | -5 | -2 | 1 | -1 | -1 | -4 | -2 | -3 | -2 | -2 | 0 |
| E | 2 | -8 | -3 | 9 | -2 | -7 | -4 | -12 | -10 | -7 | -17 | -8 | -6 | -3 | -8 | -10 | -10 | -13 | -6 | -3 | 0 |
| F | 3 | -5 | 0 | -2 | 7 | -3 | -3 | -5 | 1 | -3 | -9 | -5 | -2 | 2 | -5 | -8 | -3 | -7 | 4 | -4 | 0 |
| G | -2 | -4 | 1 | -7 | -3 | 6 | 3 | 0 | -7 | -7 | -1 | -2 | -2 | -4 | 3 | -3 | 4 | -6 | -4 | -2 | 0 |
| H | -2 | -4 | 1 | -4 | -3 | 3 | 6 | -4 | -7 | -6 | -6 | 0 | -1 | -3 | 1 | -3 | -1 | -5 | -5 | 3 | 0 |
| I | -7 | -12 | -3 | -12 | -5 | 0 | -4 | 8 | -5 | -11 | 7 | -7 | -6 | -6 | -3 | -9 | 6 | -12 | -5 | -8 | 0 |
| K | -3 | -13 | -5 | -10 | 1 | -7 | -7 | -5 | 9 | -11 | -8 | -12 | -6 | -5 | -9 | -14 | -5 | -15 | 5 | -8 | 0 |
| L | -3 | 1 | -4 | -7 | -3 | -7 | -6 | -11 | -11 | 6 | -16 | -3 | -2 | 2 | -4 | -4 | -9 | 0 | -8 | -9 | 0 |
| M | -10 | -14 | -5 | -17 | -9 | -1 | -6 | 7 | -8 | -16 | 10 | -9 | -9 | -10 | -5 | -10 | 3 | -16 | -6 | -9 | 0 |
| N | -5 | 0 | -2 | -8 | -5 | -2 | 0 | -7 | -12 | -3 | -9 | 7 | 0 | -2 | 2 | 3 | -4 | 0 | -8 | -5 | 0 |
| P | -1 | 0 | 1 | -6 | -2 | -2 | -1 | -6 | -6 | -2 | -9 | 0 | 4 | 0 | 0 | -2 | -4 | 0 | -4 | -5 | 0 |
| Q | 1 | 1 | -1 | -3 | 2 | -4 | -3 | -6 | -5 | 2 | -10 | -2 | 0 | 5 | -2 | -4 | -5 | -1 | -2 | -5 | 0 |
| R | -4 | -1 | -1 | -8 | -5 | 3 | 1 | -3 | -9 | -4 | -5 | 2 | 0 | -2 | 6 | 2 | 0 | -1 | -6 | -3 | 0 |
| S | -7 | 0 | -4 | -10 | -8 | -3 | -3 | -9 | -14 | -4 | -10 | 3 | -2 | -4 | 2 | 6 | -6 | 0 | -11 | -9 | 0 |
| T | -5 | -8 | -2 | -10 | -3 | 4 | -1 | 6 | -5 | -9 | 3 | -4 | -4 | -5 | 0 | -6 | 8 | -9 | -5 | -5 | 0 |
| V | -6 | 1 | -3 | -13 | -7 | -6 | -5 | -12 | -15 | 0 | -16 | 0 | 0 | -1 | -1 | 0 | -9 | 3 | -10 | -11 | 0 |
| W | 0 | -7 | -2 | -6 | 4 | -4 | -5 | -5 | 5 | -8 | -6 | -8 | -4 | -2 | -6 | -11 | -5 | -10 | 8 | -6 | 0 |
| Y | -2 | -9 | -2 | -3 | -4 | -2 | 3 | -8 | -8 | -9 | -9 | -5 | -5 | -5 | -3 | -9 | -5 | -11 | -6 | 9 | 0 |
| X | 0 | 0 | 0 | 0 | 0 | 0 | 0 | 0 | 0 | 0 | 0 | 0 | 0 | 0 | 0 | 0 | 0 | 0 | 0 | 0 | 0 |

The following alignment represents five consecutive columns from five aligned protein structures. Each entry corresponds to a structure-aware character assigned to a residue by Foldseek. The alignment is shown in matrix format, where each row is a sequence and each column corresponds to a structurally aligned position. This block is centered within a longer alignment and represents columns $i$ through $i$+4:

| Sequence | Col $i$ | Col $i$+1 | Col $i$+2 | Col $i$+3 | Col $i$+4 |
|---|---|---|---|---|---|
| S1 | P | V | L | A | K |
| S2 | P | V | L | A | K |
| S3 | V | P | L | Q | K |
| S4 | V | P | L | Q | K |
| S5 | D | V | Q | G | W |

Table 3: Extract from a structure-based alignment showing five columns of a longer alignment.

For each alignment column, confidence scores are computed based on pairwise substitution values between adjacent sequence rows (e.g., S1–S2, S2–S3, etc.). These scores are summed to produce a raw confidence value $C_j$ for each column. Min-max normalization is then applied across all five columns to produce normalized confidence values $\hat{C}_j$ in the range $[0, 1]$. The average of these normalized values is reported as $\hat{C}_{\text{avg}}$, representing the overall alignment confidence.

| Column | Residue Pairs | Pair Scores | $\mathbf{C_j}$ | $\mathbf{\hat{C}_j}$ |
|---|---|---|---|---|
| Col $i$ | (P,P), (P,V), (V,V), (V,D) | 4, 0, 3, -3 | 4 | 0.00 |
| Col $i$+1 | (V,V), (V,P), (P,P), (P,V) | 3, 0, 4, 0 | 7 | 0.11 |
| Col $i$+2 | (L,L), (L,L), (L,L), (L,Q) | 6, 6, 6, 2 | 20 | 0.57 |
| Col $i$+3 | (A,A), (A,Q), (Q,Q), (Q,G) | 6, 1, 5, -4 | 8 | 0.14 |
| Col $i$+4 | (K,K), (K,K), (K,K), (K,W) | 9, 9, 9, 5 | 32 | 1.00 |

Table 4: Raw and normalized confidence scores for a real alignment example across five columns. $\hat{C}_j$ is computed by min-max normalization with $C_{\text{min}} = 4$ and $C_{\text{max}} = 32$. The average normalized confidence across columns is $\hat{C}_{\text{avg}} = 0.364$.

This example highlights how confidence scores reflect local structural agreement among adjacent sequences and are scaled based on alignment-internal variation. The normalized values do not reflect absolute correctness but instead provide a relative measure to highlight which columns are most structurally consistent within the alignment.

### Supplementary Example: Alignment Statistics

The following section shows the complete alignment for SCOP family (accession d.12.1.3), based on structure-aware sequences.

```
2v94_A_1-93       DDKAWDDWDQDPVLAKIKTKIKDQDPPDDADALQVVLVCVCVVVVADSLFKFWDDWADDP
2v94_B_1-93       DDKAWDDWDQDPVLAKIKTKIKDADPPDDDDDLQVVLVCVCVVVVADSLQKFWPDWADDV
1ywx_A_1-102      DDKAWDDWDADVPLQKTKTWIKDADD-DDDDAFLVVLVVVCVVVVDDSPQWTFQDKAADP
1xn9_A            DDKDWDDKADDVPLQKIKTWIKDADD-DDADDPVRVLVVVCVVVVHDSLQKDWDDWDADD
2g1d_A_1-98       DAKDKAWAADDDVQGWTKIKIKDDDDDDDDDDLCVCQQNHVVNNVHGSQFWDDDDWDADD

2v94_A_1-93       VHRMIMTIIMGHPDPVSSVVPDDVVRSVSNVD-----------
2v94_B_1-93       VHRMIMTIIMGHPDNVSCVVPPDPVRCVSSPND----------
1ywx_A_1-102      VDRMMTGMTMGHDDVVSCVPVPDCVRCVSVDDDDDDPDDQDDD
1xn9_A            DGRMIITIIMGHPDNVSVCVSVVVVPPDDDDDDDDDDDDDDD-
2g1d_A_1-98       PHHMTIGMTITGPCRVQCVVVNPPGDPVDDDDPPVHPD-----
```

| **Statistic** | **Value** |
|---|---|
| Number of structures | 5 |
| Alignment length (columns) | 103 |
| Total positions | 515 |
| Gap fraction | 0.0563 |
| Dropped columns (>50% gaps) | 5 |
| Trimmed alignment length | 98 |
| Average confidence | 0.3512 |
| Average confidence (trimmed) | 0.3651 |

Table 5: Summary statistics for the structure-based alignment shown above.
